# KA-Search: Rapid and exhaustive sequence identity search of known antibodies

**DOI:** 10.1101/2022.11.01.513855

**Authors:** Tobias H. Olsen, Brennan Abanades, Iain H. Moal, Charlotte M. Deane

**Affiliations:** Oxford Protein Informatics Group, Department of Statistics, University of Oxford, Oxford, United Kingdom; GSK Medicines Research Centre, GlaxoSmithKline plc, Stevenage, United Kingdom; Exscientia plc, Oxford, United Kingdom

**Keywords:** antibody, sequence search, immune repertoire, immune repertoire mining, immunogenicity

## Abstract

Antibodies with similar amino acid sequences, especially across their complementary-determining regions, often share properties. Finding that an antibody of interest has a similar sequence to naturally expressed antibodies in healthy or diseased repertoires is a powerful approach for the prediction of antibody properties, such as immunogenicity or antigen specificity. However, as the number of available antibody sequences is now in the billions and continuing to grow, repertoire mining for similar sequences has become increasingly computationally expensive. Existing approaches are limited by either being low-throughput, non-exhaustive, not antibody specific, or only searching against entire chain sequences. Therefore, there is a need for a specialized tool, optimized for a rapid and exhaustive search of any antibody region against all known antibodies, to better utilize the full breadth of available repertoire sequences.

We introduce Known Antibody Search (KA-Search), a tool that allows for the rapid search of billions of antibody sequences by sequence identity across either the whole chain, the complementarity-determining regions, or a user defined antibody region. We show KA-Search in operation on the ∼2.4 billion antibody sequences available in the OAS database. KA-Search can be used to find the most similar sequences from OAS within 30 minutes using 5 CPUs. We give examples of how KA-Search can be used to obtain new insights about an antibody of interest. KA-Search is freely available at https://github.com/oxpig/kasearch.

## 1 INTRODUCTION

Antibodies have become an invaluable form of therapeutics, with an increasing number of new antibody derived therapeutics being developed and marketed each year (1). Despite their success, the process of antibody discovery and design is still challenging (2). One complication is the huge mutational space of antibodies that has to be explored in order to find the desired binding specificity alongside other properties. A technique which has shown promise for exploring the mutational space of antibodies is immune repertoire mining (3, 4, 5, 6). In immune repertoire mining, an antibody of interest is compared against natural antibody repertoires to find identical or highly similar antibodies. This is useful, as similar antibodies often share properties and it can therefore be a powerful method for finding antibodies in nature which have improved properties such as their developability profile, reduced immunogenicity or increased affinity (3, 4, 5, 6, 7) (see Figure 1A).

**Figure 1.**
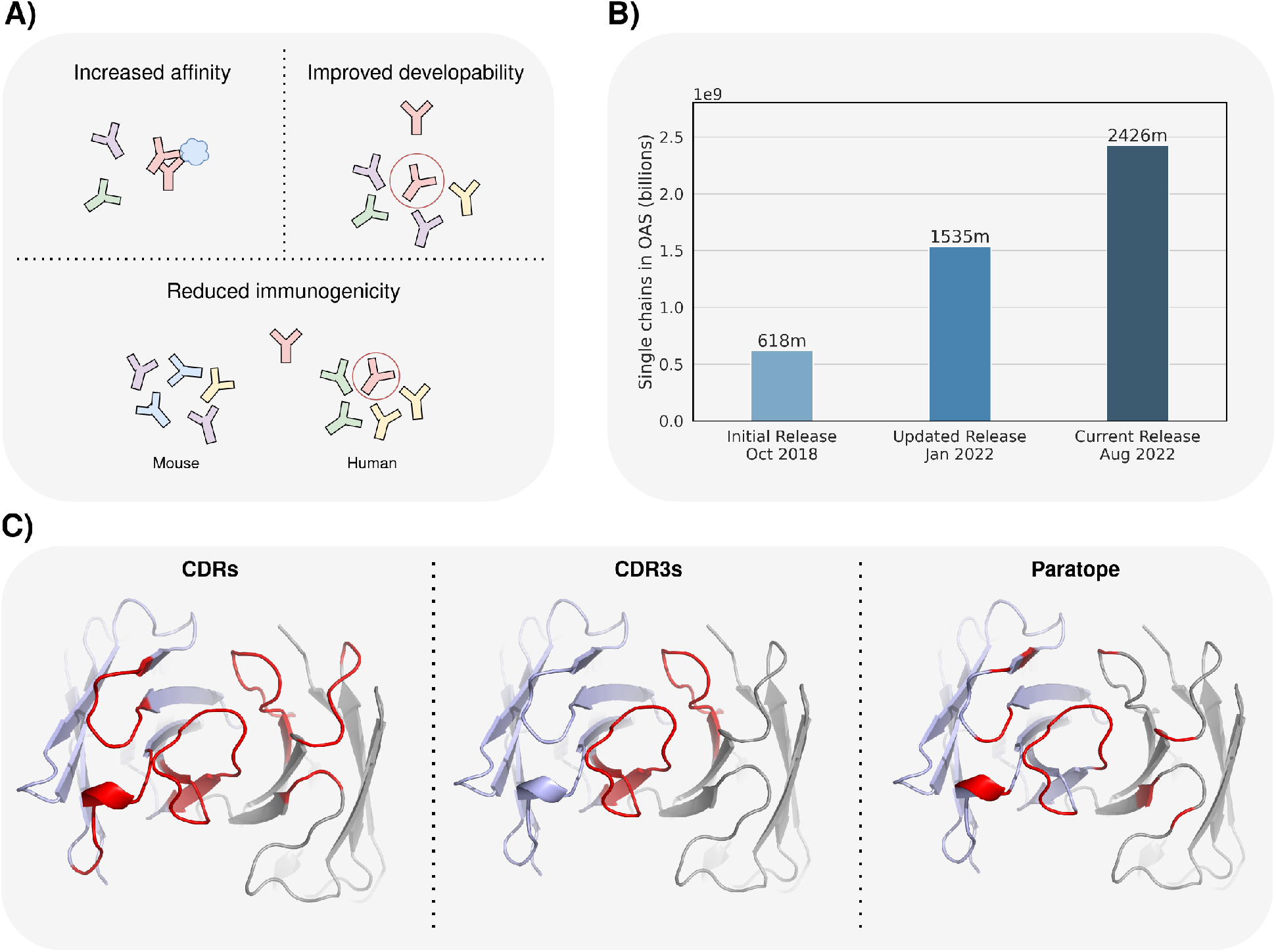
A) Similar antibodies often share properties. Immune repertoire mining can therefore be used to find similar binders with an increased affinity, an improved developability profile or reduced immunogenicity. B) Overview of the available number of single chain antibody sequences in the Observed Antibody Space database over time. C) Highlight (red) of different specific search regions, useful for finding similar binders. Complementary-determining region (CDR).

Similarity between antibodies can be measured in different ways. The most common ones are via sequence identity or structural similarity (8, 9). With a protein’s function being preserved in the structure, structural similarity is often superior for finding proteins with analogous functions, such as antibodies binding the same epitope (7, 10). However, with orders of magnitude more sequence data available than structural data, a sequence identity search enables the exploration of a much larger space. Sequence data is also more diverse, as next generation sequencing of B-cell receptors (BCR) is routinely being applied to study adaptive immunity, generating sequences from a range of species (11, 12, 13) and from individuals with differing disease states (14, 15). Furthermore, continuous improvements in high-throughput sequencing methods and increased adoption by research labs means that the amount and diversity of sequence data is rapidly increasing (16, 17).

Searching all this immune repertoire data for similar sequences is useful for a wide range of applications, such as finding the most similar human antibody sequence to an antibody isolated from an animal model during therapeutic development. While freely available, searching this data requires extensive post-processing of each source, and a database providing a single entry to antibody data to search against is therefore advantageous. One such effort is the Observed Antibody Space (OAS) (18, 19) database, which collates data from publicly available BCR sequencing studies and as of September 2022 contains ∼2.4 billion unpaired heavy and light antibody chains. While the size of OAS is promising from a scientific perspective, its scale and continuous growth, visualized in Figure 1B, make effectively mining it a challenge. Calculating the sequence identity between antibodies is simple, but without software specially optimised for the task, the computational cost of exhaustively searching OAS and other large antibody sequence databases is becoming prohibitive. There is therefore a need for specialized tools to search this space now and in the future.

There exist many tools for searching large datasets of protein sequences for similar sequences, for example BLASTp (20) and CD-Hit-2d (21), and newer methods such as MMseqs2 (22). However, these tools are designed around searching a diverse set of proteins and often exploit the low similarity between most sequences. To increase speed, both CD-Hit and MMseqs2 prefilter the target sequences for low identity sequences to reduce the number of pairwise alignments to make, as this is a computational expensive step (22). However, this is not as effective for closely related sequences such as antibodies, as the prefiltering can remove good hits. Further, each tool uses an alignment method designed for general protein sequences, which can result in unreliable antibody alignments, especially in the highly variable complementary-determining regions (CDRs). Within the immunoinformatics field this alignment problem is often overcome by using antibody specific numbering schemes, like the ImMunoGeneTics (IMGT) scheme (23). Another issue with non-antibody specific tools, is the lack of flexibility in their searches. These tools can only readily be used on the whole antibody chain and not for finding similar sequences based on subregions. Searching for specific, identical regions within antibodies, especially the CDRs, is often used when looking for similar binders (6). With the majority of the residues involved in binding being located in the CDRs, the sequence identity over this region is often more relevant than that of the whole antibody. For some applications, the exact set of residues involved in binding (paratope) may be known. In these cases, searching based on the sequence identity of the paratope may be even more informative (see Figure 1C). An antibody specific tool utilizing antibody numbering schemes for better searches, without prefiltering for an exhaustive search, and with the ability to search user-defined regions would improve our ability to make best use of the antibody sequence data available.

Recent efforts to create antibody specific searching tools include iReceptor (24) and AbDiver (25). iReceptor, only allows for a V-, D-, or J-gene search or an exact CDR3 match search. AbDiver uses an antibody numbering scheme to align sequences and allows for both CDR3 and whole chain searches. AbDiver restricts CDR3 searches against CDR3s with a specified V gene and species of origin, and whole chain searches against sequences with same length CDR1 and 2 and *±*1 length CDR3. These restrictions narrow and greatly speed up the search but can occasionally lead to it finding no matches. Further, both iReceptor and AbDiver are not open-source and are only freely available to use via their website, so can only be used against their own databases. There therefore exists the need for an open-source antibody specific tool not limited by either being low-throughput, non-exhaustive, or only searching against entire chain sequences.

Here, we introduce Known Antibody Search (KA-Search), a tool that allows for rapid sequence identity search across billions of unpaired antibody chains, across either the whole chain, the CDRs, or a user defined antibody region. We demonstrate KA-Search can be used to find the most similar sequences from the ∼ 2 billion heavy chain sequences in the OAS database within 30 minutes using 5 CPUs. We also show how KA-Search can be used for immune repertoire mining to obtain new insights about an antibody of interest. KA-Search is freely available at https://github.com/oxpig/kasearch.

## 2 METHOD

### 2.1 Data preprocessing

KA-Search pre-aligns antibody sequences to a canonical alignment capable of accommodating the most common numbering positions. To do this, every antibody sequence is first numbered with ANARCI (26) using the IMGT numbering scheme (27) and then converted to a vector of the same length to enable fast matrix calculations. As our canonical alignment we use a set of 196 unique positions seen in at least 40,000 different sequences in OAS, as of May 2022, and four additional unique positions seen in therapeutics from Thera-SAbDab (28). The exact unique positions are given in Table S1 and cover around ∼99.8% of sequences in OAS. The 0.2% of antibody sequences that contain a rare insertion can not be aligned and are hence compared using a slower method. Every aligned sequence is accompanied by two index values which can be used to retrieve its meta data.

All sequences in OAS (September 2022) are pre-aligned using this method to generate a dataset ready to be used by KA-Search. This results in over 2,070 million heavy and 355 million light chain sequences. Sequences are split into heavy and light chains, and by species information, e.g. human, mouse, rabbit, rat, rhesus, camel and humanized, allowing for faster specific searches. We call this data set of heavy and light chains for OAS-aligned. We also built a subset of the heavy chain dataset, OAS-aligned-small, that contains 144 million human heavy chain sequences. This was generated by removing sequences containing ambiguous residues or seen less than three times. OAS-aligned and OAS-aligned-small, and the code to update the data sets or expand it with an in-house data set is made freely available with KA-Search (https://github.com/oxpig/kasearch).

### 2.2 Identity calculation

The identity between the query sequence and a target sequence is computed using the method as described by Krawczyk et al., 2019 (8) for both length matched and not length matched. To search for the identity of a specific user-defined region, i.e. the CDRs, a list with the desired positions can be specified. These positions need to be one of the 200 unique positions in the canonical alignment. In default mode, KA-Search will search for similar whole chains of variable length, and the three CDRs and CDR3 regions with exact length match.

### 2.3 Sensitivity and speed comparison

KA-Search was compared to BLASTp (version 2.13.0), CD-Hit-2d (version 4.8.1) and MMseqs2 (version 13.45111), for sensitivity and speed at searching for the closest whole antibody chain match. A set of 100 randomly selected non-redundant heavy chains of therapeutics were used to search for the closest sequence within OAS-test, a set of 10 million heavy chain antibody sequences. The sequences in OAS-test were randomly extracted from the full OAS database, cleaned and reduced as done in (29). The 100 therapeutic heavy chains and OAS-test are available for download, see Data Availability Statement.

For each tool we used the default settings. For BLASTp we searched against a pre-built BLAST database of OAS-test, for MMseqs2 we used the easy-search workflow to search a pre-computed sequence database of OAS-test, and for KA-Search we used a pre-aligned OAS-test, aligned as described above. Speed was calculated using the same single CPU to search for one sequence against OAS-test and sensitivity by how well each tool found the closest sequence in OAS-test to each query. As KA-Search is an exhaustive method, it provides the ground truth for the sensitivity comparison.

## 3 RESULTS

Immune repertoire mining to find similar antibodies with shared properties is becoming increasingly computational expensive because of the increase in available antibody sequences. This is illustrated in Figure 1B, which shows how publicly available sequences in OAS have increased by 1.8 billion in less than four years. Below we describe KA-Search, a freely available tool to search immune repertoires that is optimised to handle the vast amount of available data.

### 3.1 Computational speed of KA-Search

KA-Search’s exact speed is dependent on the hardware used, the number of queries, number of output sequences desired and number of regions searched over. Figure 2 shows a comparison between different KA-Search runs with different numbers of CPUs, when searching with a single antibody heavy chain against the 2,070 million heavy chains in OAS-aligned. The number of closest matches returned has a minimal impact on speed, with returning the best or 10,000 best matches taking approximately the same time. Searching over multiple regions simultaneously slows the search but is faster than doing them individually. When using a single CPU one region takes ∼42, three ∼52, and ten ∼127 minutes. The time required per query is reduced when searching with multiple queries at a time, as searching with a single query takes ∼42 minutes while searching with 100 queries takes ∼4.3 minutes per query. KA-Search is limited by loading data into memory when searching with few queries. The optimal use of KA-Search is therefore to search with many queries and multiple regions simultaneously using multiple CPUs. All time measurements in this paper were performed using CPUs from an Intel Xeon Gold 6240 Processor.

**Figure 2.**
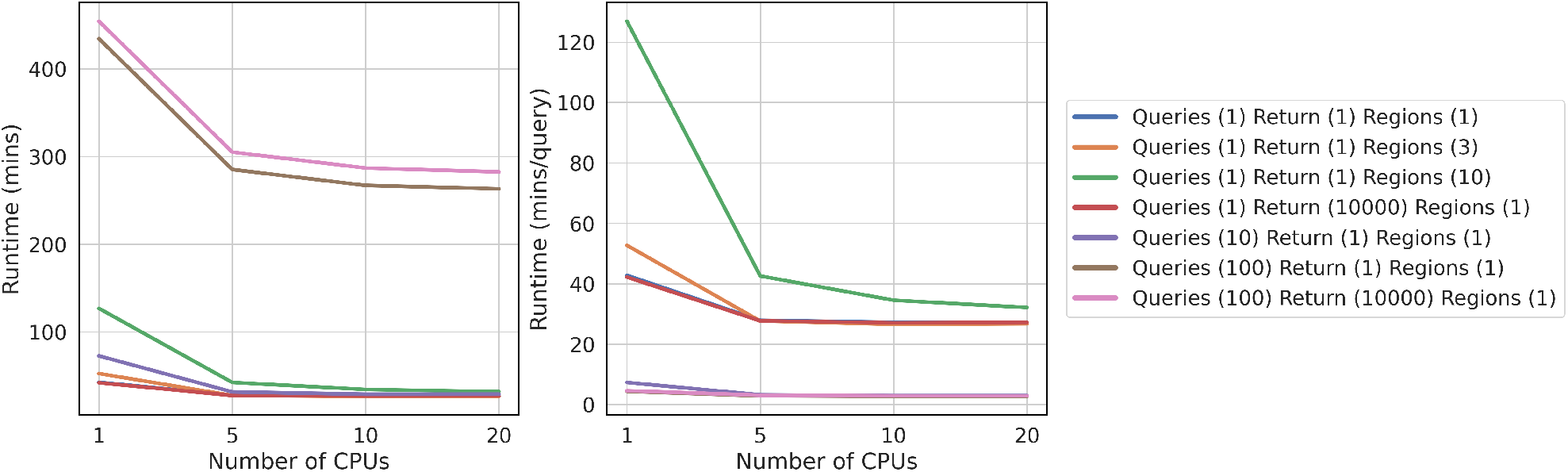
Running KA-Search with different numbers of queries, returned sequences and number of regions, when searching with a single antibody heavy chain against the 2,070 million heavy chains in the Observed Antibody Space database.

### 3.2 Sensitivity comparison

To compare KA-Search with current freely available and downloadable protein sequence search tools, we selected 100 non-redundant heavy chains of therapeutics, and searched for the most similar sequence within OAS-test, a set of 10 million heavy chain antibody sequences (see methods), using BLASTp (20), CD-Hit-2d (21), MMseqs2 (22) and KA-Search (Figure 3). KA-Search takes ∼8 seconds, which is far faster than BLASTp and CD-Hit-2d, ∼103 and ∼82 seconds respectively, but slower than MMseqs2 at ∼3 seconds. In terms of sensitivity, identifying the closest antibody in OAS-test, the conventional tools struggle to find the exact closest match, with BLASTp finding the exact match as highest ranked for 19 out of the 100 sequences, CD-Hit-2d for three sequences and MMseqs2 for no sequences. When looking for the closest match within the top-100 highest ranked sequences, BLASTp and CD-Hit-2d find the closest match for 65 and 14 sequences, respectively, while MMseqs2 finds none. The average difference in identity between the highest ranked and closest match was also calculated. For BLASTp, CD-Hit-2d and MMseqs2 this difference was on average 3.93%, 11.34% and 7.15% identity, respectively. The difference between the closest match within the 10 million sequences and the best within the top-100 highest ranked sequences were 0.27%, 4.53% and 4.0%, respectively. While CD-Hit-2d is better than MMseqs2 at finding the closest match, MMseqs2 returns on average better matches. As KA-Search is exhaustive, it finds the exact closest match every time.

**Figure 3.**
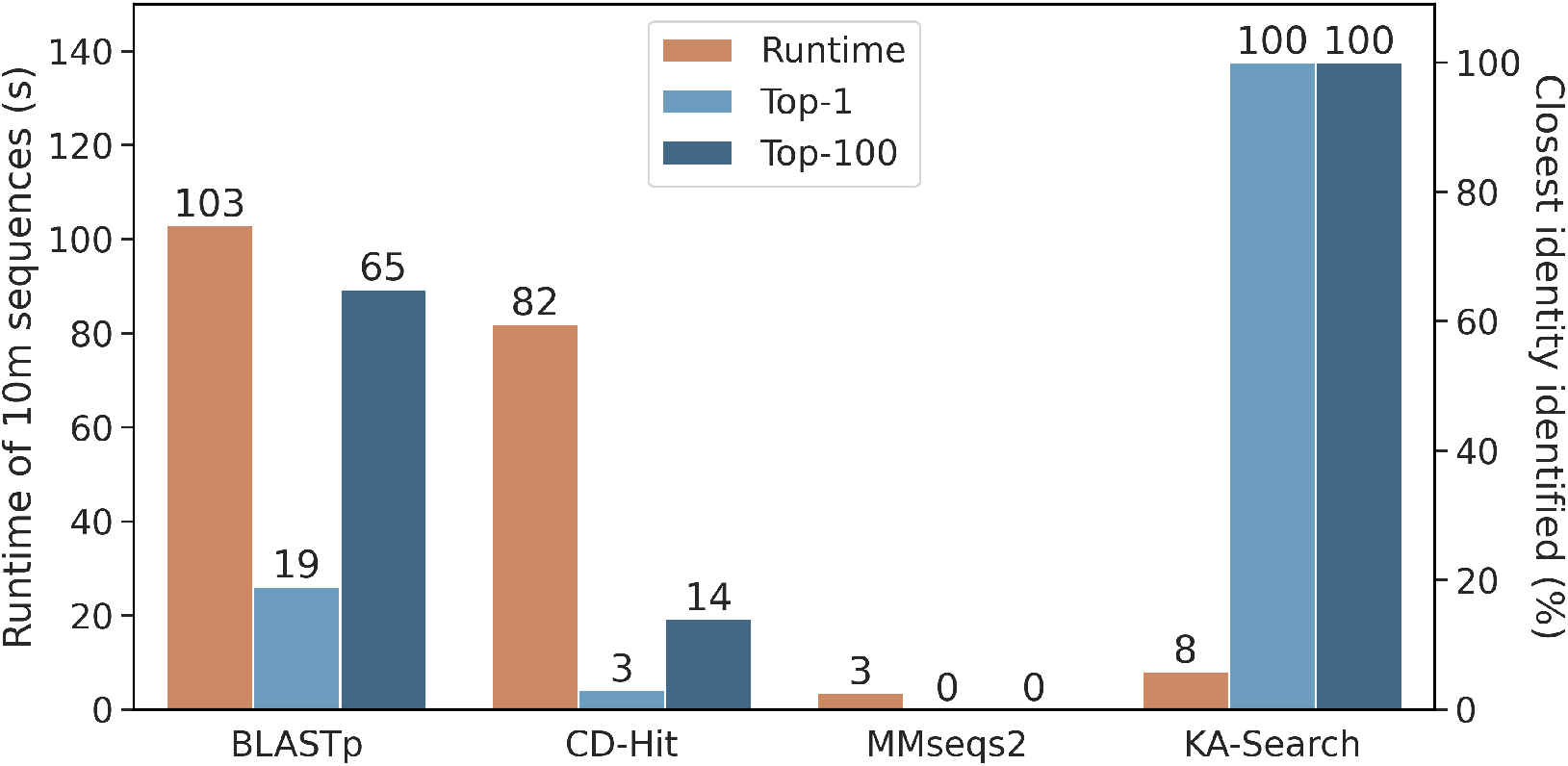
Speed and sensitivity comparison between KA-Search and the commonly used protein search tools BLASTp (version 2.13.0), CD-Hit-2d (version 4.8.1) and MMseqs2 (version 13.45111). The default setting was used for each tool. Speed was calculated as the time it took to search 10 million sequences (OAS-test) with a single query on a single CPU and sensitivity by how often the tools returned the closest or the closest within the top-100 match for 100 heavy chains against the same 10 million sequences. As KA-Search is an exhaustive method, it provides the ground truth for the sensitivity comparison.

### 3.3 Immune repertoire mining with the COVOX-253 antibody

COVOX-253 is an antibody which binds to the neck of SARS-CoV-2’s Receptor-Binding Domain (RBD) (30). Using KA-Search, up to the 1,000 closest sequences to the heavy chain of COVOX-253, with over 90% identity, were extracted for four different regions: the whole antibody chain, the three CDRs, the CDR3 and the paratope. The paratope was derived from the PDB structure 7BEN and defined as any residue in the antibody which was within 4.5Å of the RBD. Figure 4A shows the disease of the patient the antibody sequence found in OAS comes from and in Figure 4B each antibody’s combination of V and J genes. Most matched sequences are derived from the genes IGHV1-58*01 and IGHJ3*02, however, COVOX-253’s CDR3 is seen with six different V genes, IGHV1-58*01, IGHV1-58*02, IGHV1-18*01, IGHV1-46*01, IGHV1-69*10 and IGHV1-69*13.

**Figure 4.**
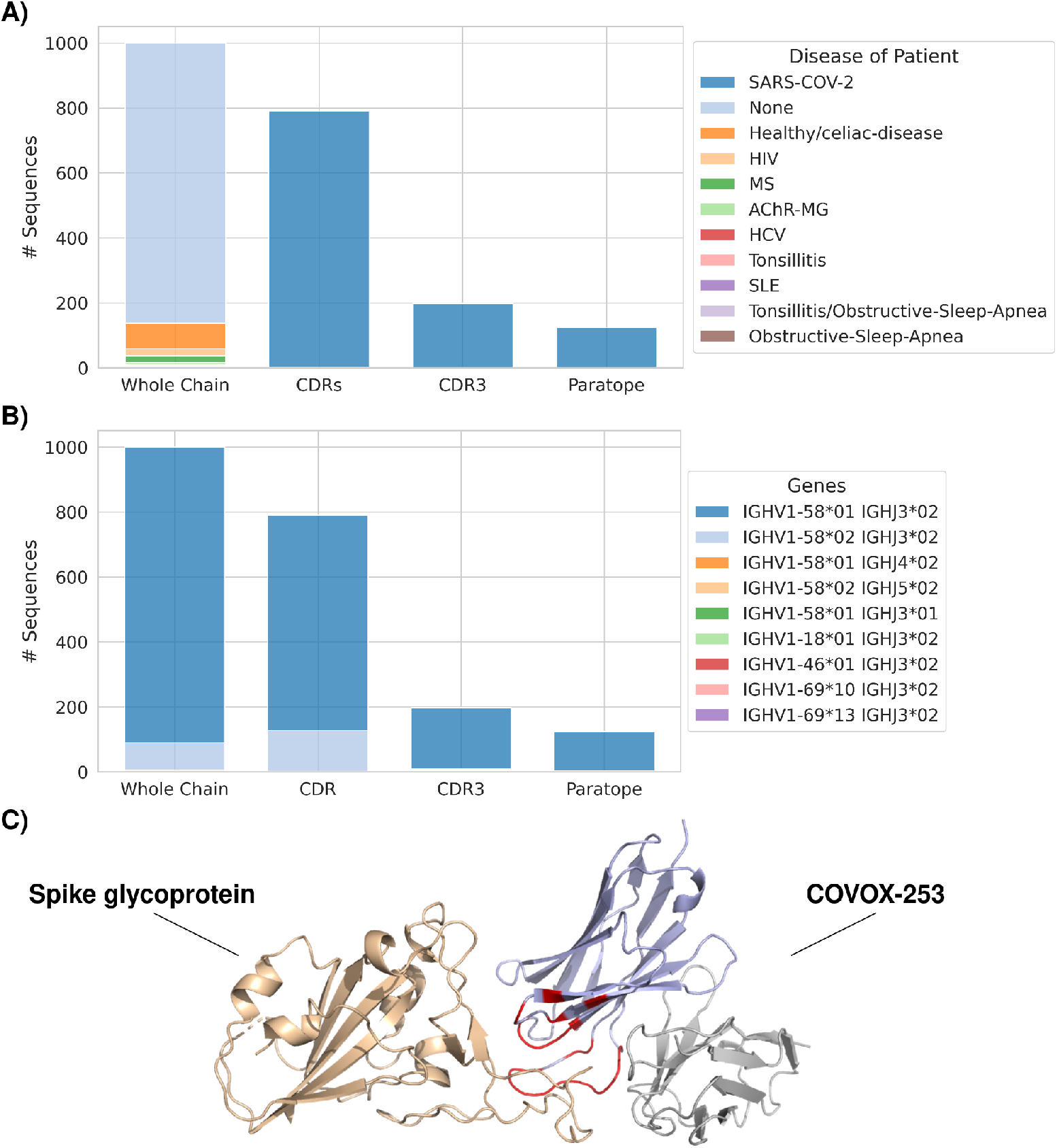
A KA-Search of the heavy chain of the SARS-CoV-2 RBD binding antibody COVOX-253. Returned antibodies with over 90% identity across four different regions were visualised based on, A) the disease state of the patient and B) which V- and J-genes the antibody sequence is derived from. C) The variable region of COVOX-253’s heavy (purple) and light (grey) chain with the bound spike glycoprotein (beige), derived from PDB 7BEN. The paratope of the heavy chain, which was used to search with KA-Search, is shown in red.

Searching for the closest match using the whole chain returns 863 sequences from healthy individuals and 137 from patients with one of seven different diseases, none which are SARS-CoV-2. Searching with the CDR positions returns one sequence from a healthy individual and 789 sequences from patients with SARS-CoV-2, while searching with the CDR3 or paratope positions returns 197 and 124 sequences, respectively, all from SARS-CoV-2 infected patients. The fact that OAS-aligned only contains ∼84 million heavy chains from patients with SARS-CoV-2, ∼4% of all heavy chains in OAS-aligned, highlights the importance of being able to search over specific regions.

## 4 DISCUSSION

Immune repertoire mining is a powerful method for identifying antibodies in nature which are similar to an antibody of interest and can help indicate likely specificity or immunogenicity. However, the number of available antibody sequences are now in the billions and is continuing to grow. Therefore, repertoire mining for highly similar sequences has become increasingly computationally expensive. Existing approaches are limited by either being low-throughput, inaccessible for large scale searches, non-exhaustive, not antibody specific, or only searching against entire chain sequences. There is therefore a need for a specialized tool, optimized for a rapid and exhaustive search of any antibody region against all known antibodies, to better utilize the full number of available repertoire sequences.

In this paper, we introduce Known Antibody Search (KA-Search), a platform independent antibody search tool. KA-Search exploits antibody numbering, allowing us to pre-align antibody sequences to a fixed-length vector. This circumvents pairwise alignment during search, an otherwise time-consuming step, and generates better alignments than tools using generic methods. The increased speed allows KA-Search to avoid prefiltering and be exhaustive while still retaining a competitive speed. Avoiding prefiltering is crucial, as current prefiltering techniques greatly reduce sensitivity when searching highly related proteins, such as antibodies, where a single mutation can be of great importance. While pre-aligning the antibody sequences increases search speed, the initial pre-alignment is slow. We therefore provide a pre-aligned dataset of the current OAS, ready to use for searching. This dataset can be extended with future OAS updates or in-house data without the need to re-align the existing sequences.

Pre-aligning sequences also opens new use-cases. Instead of only searching against the whole antibody chain, searches can now be focused on specific positions in the alignment. Searches can be specific for the CDRs or regions specific for individual antibodies, such as the paratope. This flexibility allows for studies which were previously difficult to execute. As an example, previously an extensive study was needed to search OAS across the whole chain, CDRs and CDR3 for the closest match to a set of 242 therapeutics (8). The same study can now be done on the 804 therapeutics within Thera-SAbDab (28), August 2022, with KA-Search in less than two days compute and little configuration (see Figure S1). KA-Search can also extract the meta data from OAS for the matched sequences, which can be used to obtain new insights about an antibody of interest. Using KA-Search to find the closest sequences with or above 90% identity across four different regions for the SARS-CoV-2 RBD binding COVOX-253 demonstrates the power of searching across particular regions. The closest matches from searching with the whole chain comes from healthy patients or patients with a variety of diseases. However, binding region specific searches only return sequences found in SARS-CoV-2 infected patients. These sequences could therefore also have RBD binding properties and are possible candidates for further affinity studies. Investigating the genes of the closest sequences, also potentially indicates which other frameworks a region of interest could exist on. For COVOX-253, the CDR3 is seen in sequences with six different V genes, which each could be possible framework candidates for the CDR3. The ability to return high numbers of close matches without decreasing speed, also opens up KA-Search as a means for creating multiple sequence alignments of similar antibody sequences.

KA-Search’s speed, exhaustiveness and flexibility allows it to search the vast numbers of antibody sequences now available seamlessly, find viable mutations and gain new insight into antibodies of interest. We therefore believe KA-Search is a useful tool that will allow the antibody community to explore antibodies in new ways. To maximize KA-Search’s possible contribution to the community, KA-Search is open source and freely available at https://github.com/oxpig/kasearch.

## AUTHOR CONTRIBUTIONS

TO, BA, IM and CD contributed to conception and design of the study. TO organised the datasets. TO and BA wrote the code and created the package. TO and BA wrote the first draft of the manuscript. All authors contributed to manuscript revision, read, and approved the submitted version.

## ACKNOWLEDGMENTS AND FUNDING

This work was supported by the Engineering and Physical Sciences Research Council [EP/S024093/1], GlaxoSmithKline Pharmaceuticals Ltd and F. Hoffmann-La Roche Ltd.

For the purpose of Open Access, the author has applied a CC BY public copyright licence to any Author Accepted Manuscript (AAM) version arising from this submission.

## SUPPLEMENTAL DATA

### DATA AVAILABILITY STATEMENT

The data sets generated for the sensitivity comparison in this study can be found in the following Zenodo repository https://doi.org/10.5281/zenodo.7190267. Links for the OAS-aligned and OAS-aligned-small can be found on the associated github https://github.com/oxpig/kasearch.

## REFERENCES

1. Kaplon H, Chenoweth A, Crescioli S, Reichert JM. Antibodies to watch in 2022. mAbs (2022), 14 2014296. doi:10.1080/19420862.2021.2014296.

2. Raybould MIJ, Marks C, Krawczyk K, Taddese B, Nowak J, Lewis AP, et al. Five computational developability guidelines for therapeutic antibody profiling. Proceedings of the National Academy of Sciences (2019), 116 4025–4030. doi:10.1073/pnas.1810576116.

3. Wang B, Kluwe CA, Lungu OI, DeKosky BJ, Kerr SA, Johnson EL, et al. Facile Discovery of a Diverse Panel of Anti-Ebola Virus Antibodies by Immune Repertoire Mining. Scientific Reports (2015), 5 13926. doi:10.1038/srep13926.

4. Tian X, Li C, Wu Y, Ying T. Deep Mining of Human Antibody Repertoires: Concepts, Methodologies, and Applications. Small Methods (2020), 4 2000451. doi:https://doi.org/10.1002/smtd.202000451.

5. Hsiao YC, Shang Y, DiCara DM, Yee A, Lai J, Kim SH, et al. Immune repertoire mining for rapid affinity optimization of mouse monoclonal antibodies. mAbs (2019), 11 735–746. doi:10.1080/19420862.2019.1584517.

6. Richardson E, Galson JD, Kellam P, Kelly DF, Smith SE, Palser A, et al. A computational method for immune repertoire mining that identifies novel binders from different clonotypes, demonstrated by identifying anti-pertussis toxoid antibodies. mAbs (2021), 13 1869406. doi:10.1080/19420862.2020.1869406.

7. Robinson SA, Raybould MIJ, Schneider C, Wong WK, Marks C, Deane CM. Epitope profiling using computational structural modelling demonstrated on coronavirus-binding antibodies. PLOS Computational Biology (2021), 17 1–20. doi:10.1371/journal.pcbi.1009675.

8. Krawczyk K, Raybould MIJ, Kovaltsuk A, Deane CM. Looking for therapeutic antibodies in next-generation sequencing repositories. mAbs (2019), 11 1197–1205. doi:10.1080/19420862.2019.1633884.

9. Krawczyk K, Kelm S, Kovaltsuk A, Galson JD, Kelly D, Trück J, et al. Structurally Mapping Antibody Repertoires. Frontiers in immunology (2018), 9 1698. doi:10.3389/fimmu.2018.01698.

10. van Kempen M, Kim SS, Tumescheit C, Mirdita M, Gilchrist CLM, Söding J, et al. Foldseek: fast and accurate protein structure search. bioRxiv (2022) 2022.02.07.479398. doi:10.1101/2022.02.07.479398.

11. Li X, Duan X, Yang K, Zhang W, Zhang C, Fu L, et al. Comparative Analysis of Immune Repertoires between Bactrian Camel’s Conventional and Heavy-Chain Antibodies. PLOS ONE (2016), 11 e0161801.

12. Corcoran MM, Phad GE, Bernat NV, Stahl-Hennig C, Sumida N, Persson MAA, et al. Production of individualized V gene databases reveals high levels of immunoglobulin genetic diversity. Nature Communications (2016), 7 13642. doi:10.1038/ncomms13642.

13. Cui A, Di Niro R, Vander Heiden JA, Briggs AW, Adams K, Gilbert T, et al. A Model of Somatic Hypermutation Targeting in Mice Based on High-Throughput Ig Sequencing Data. The Journal of Immunology (2016), 197 3566LP–3574. doi:10.4049/jimmunol.1502263.

14. Johnson EL, Doria-Rose NA, Gorman J, Bhiman JN, Schramm CA, Vu AQ, et al. Sequencing HIV-neutralizing antibody exons and introns reveals detailed aspects of lineage maturation. Nature Communications (2018), 9 4136. doi:10.1038/s41467-018-06424-6.

15. Bernardes JP, Mishra N, Tran F, Bahmer T, Best L, Blase JI, et al. Longitudinal Multi-omics Analyses Identify Responses of Megakaryocytes, Erythroid Cells, and Plasmablasts as Hallmarks of Severe COVID-19. Immunity (2020), 53 1296–1314. doi:10.1016/j.immuni.2020.11.017.

16. Briney B, Inderbitzin A, Joyce C, Burton DR. Commonality despite exceptional diversity in the baseline human antibody repertoire. Nature (2019), 566 393–397. doi:10.1038/s41586-019-0879-y.

17. Soto C, Bombardi RG, Branchizio A, Kose N, Matta P, Sevy AM, et al. High frequency of shared clonotypes in human B cell receptor repertoires. Nature (2019), 566 398–402. doi:10.1038/s41586-019-0934-8.

18. Kovaltsuk A, Leem J, Kelm S, Snowden J, Deane CM, Krawczyk K. Observed Antibody Space: A Resource for Data Mining Next-Generation Sequencing of Antibody Repertoires. The Journal of Immunology (2018), 201 2502–2509. doi:10.4049/jimmunol.1800708.

19. Olsen TH, Boyles F, Deane CM. OAS: A diverse database of cleaned, annotated and translated unpaired and paired antibody sequences. Protein science: a publication of the Protein Society (2021). doi:10.1002/pro.4205.

20. Camacho C, Coulouris G, Avagyan V, Ma N, Papadopoulos J, Bealer K, et al. BLAST+: architecture and applications. BMC Bioinformatics (2009), 10 421. doi:10.1186/1471-2105-10-421.

21. Fu L, Niu B, Zhu Z, Wu S, Li W. CD-HIT: accelerated for clustering the next-generation sequencing data. Bioinformatics (Oxford, England) (2012), 28 3150–3152. doi:10.1093/bioinformatics/bts565.

22. Steinegger M, Söding J. Clustering huge protein sequence sets in linear time. Nature Communications (2018), 9 2542. doi:10.1038/s41467-018-04964-5.

23. Giudicelli V, Chaume D, Lefranc MP. IMGT/GENE-DB: a comprehensive database for human and mouse immunoglobulin and T cell receptor genes. Nucleic acids research (2005), 33 256–61. doi:10.1093/nar/gki010.

24. Corrie BD, Marthandan N, Zimonja B, Jaglale J, Zhou Y, Barr E, et al. iReceptor: A platform for querying and analyzing antibody/B-cell and T-cell receptor repertoire data across federated repositories. Immunological reviews (2018), 284 24–41. doi:10.1111/imr.12666.

25. Młokosiewicz J, Deszyński P, Wilman W, Jaszczyszyn I, Ganesan R, Kovaltsuk A, et al. AbDiver: a tool to explore the natural antibody landscape to aid therapeutic design. Bioinformatics (2022), 38 2628–2630. doi:10.1093/bioinformatics/btac151.

26. Dunbar J, Deane CM. ANARCI: antigen receptor numbering and receptor classification. Bioinformatics (Oxford, England) (2016), 32 298–300. doi:10.1093/bioinformatics/btv552.

27. Giudicelli V, Brochet X, Lefranc MP. IMGT/V-QUEST: IMGT standardized analysis of the immunoglobulin (IG) and T cell receptor (TR) nucleotide sequences. Cold Spring Harbor protocols (2011), 2011 695–715. doi:10.1101/pdb.prot5633.

28. Raybould MIJ, Marks C, Lewis AP, Shi J, Bujotzek A, Taddese B, et al. Thera-SAbDab: the Therapeutic Structural Antibody Database. Nucleic Acids Research (2020), 48 D383–D388. doi:10.1093/nar/gkz827.

29. Olsen TH, Moal IH, Deane CM. AbLang: an antibody language model for completing antibody sequences. Bioinformatics Advances (2022), 2 vbac046. doi:10.1093/bioadv/vbac046.

30. Dejnirattisai W, Zhou D, Ginn HM, Duyvesteyn HME, Supasa P, Case JB, et al. The antigenic anatomy of SARS-CoV-2 receptor binding domain. Cell (2021), 184 2183–2200. doi:10.1016/j.cell.2021.02.032.

