## Supplementary material for "KA-Search: Rapid and exhaustive sequence identity search of known antibodies": Table S1

### 1 THE CANONICAL ALIGNMENT

**Table S1.** The 200 unique IMGT (Lefranc et al., 2015) positions in our canonical alignment. We choose 196 unique positions seen in at least 40.000 different sequences in OAS, as of May 2022, and four additional unique positions seen in therapeutics from Thera-SAbDab (Raybould et al., 2020). The four additional positions are 3A, 51A, 85C and 85D.

| | The canonical alignment's unique IMGT positions |
|---|---|
| **FRW1** | 1, 2, 3, 3A, 4, 5, 6, 7, 8, 9, 10, 11, 12, 13, 14, 15, 16, 17, 18, 19, 20, 21, 22, 23, 24, 25, 26 |
| **CDR1** | 27, 28, 29, 30, 31, 32, 32A, 32B, 33C, 33B, 33A, 33, 34, 35, 36, 37, 38 |
| **FRW2** | 39, 40, 40A, 41, 42, 43, 44, 44A, 45, 45A, 46, 46A, 47, 47A, 48, 48A, 48B, 49, 49A, 50, 51, 51A, 52, 53, 54, 55 |
| **CDR2** | 56, 57, 58, 59, 60, 60A, 60B, 60C, 60D, 61E, 61D, 61C, 61B, 61A, 61, 62, 63, 64, 65 |
| **FRW3** | 66, 67, 67A, 67B, 68, 68A, 68B, 69, 69A, 69B, 70, 71, 71A, 71B, 72, 73, 73A, 73B, 74, 75, 76, 77, 78, 79, 80, 80A, 81, 81A, 81B, 81C, 82, 82A, 83, 83A, 83B, 84, 85, 85A, 85B, 85C, 85D, 86, 86A, 87, 88, 89, 90, 91, 92, 93, 94, 95, 96, 96A, 97, 98, 99, 100, 101, 102, 103, 104 |
| **CDR3** | 105, 106, 107, 108, 109, 110, 111, 111A, 111B, 111C, 111D, 111E, 111F, 111G, 111H, 111I, 111J, 111K, 111L, 112L, 112K, 112J, 112I, 112H, 112G, 112F, 112E, 112D, 112C, 112B, 112A, 112, 113, 114, 115, 116, 117, 118 |
| **FRW4** | 119, 119A, 120, 121, 122, 123, 124, 125, 126, 127, 128 |

### 2 CLOSEST MATCHES TO THERAPEUTIC ANTIBODIES

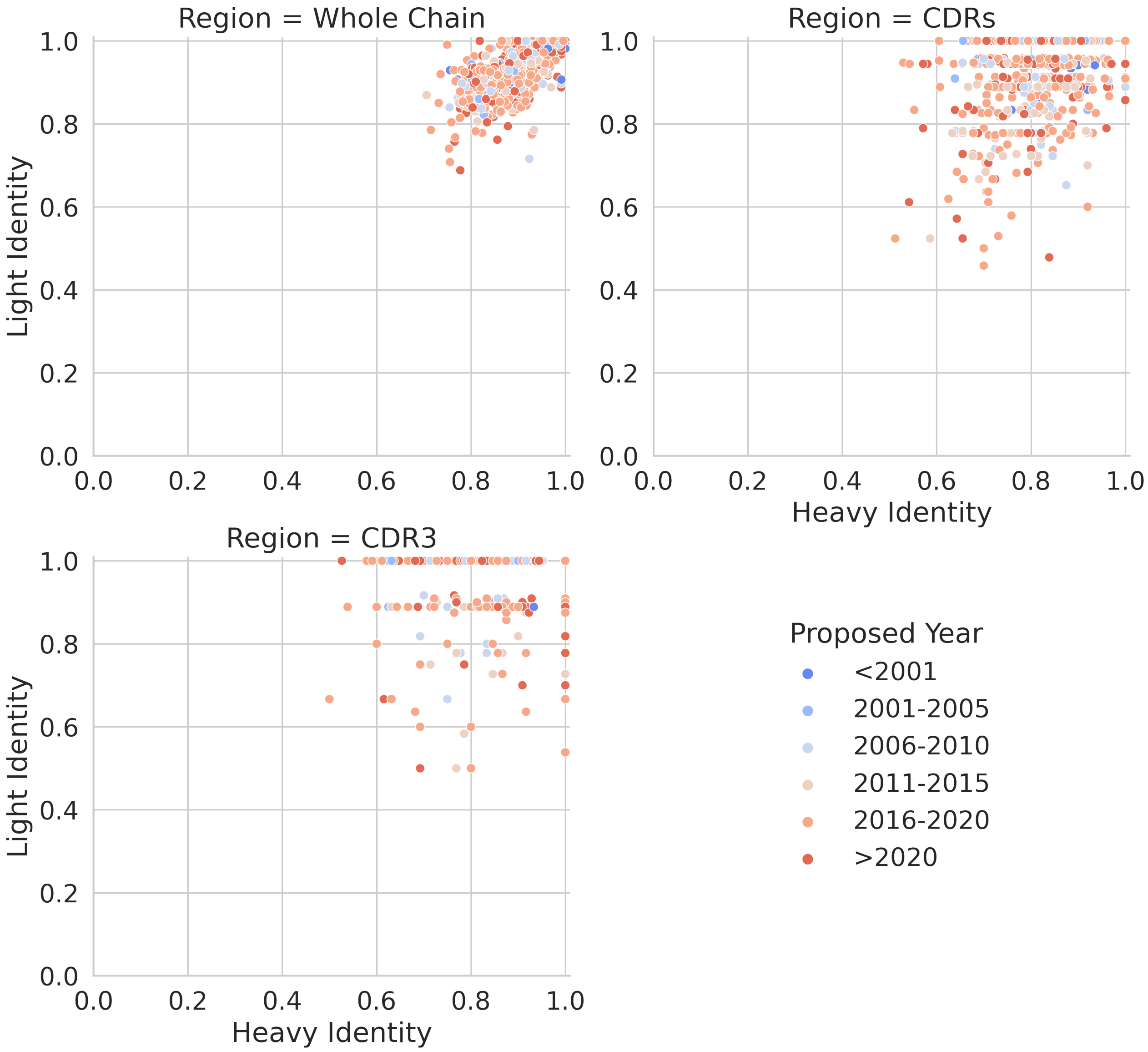


**Figure S1.** KA-Search was used to find the closest matches in OAS to 804 therapeutics extracted in August 2022 from Thera-SAbDab (Raybould et al., 2020). Closest matches was found across the the whole chain, the three CDRs and the CDR3. Each point is colored by the year they were proposed.

### REFERENCES

Lefranc MP, Giudicelli V, Duroux P, Jabado-Michaloud J, Folch G, Aouinti S, et al. IMGT®, the international ImMunoGeneTics information system® 25 years on. *Nucleic Acids Research* (2015), **43** D413–D422. doi:10.1093/nar/gku1056.

Raybould MIJ, Marks C, Lewis AP, Shi J, Bujotzek A, Taddese B, et al. Thera-SAbDab: the Therapeutic Structural Antibody Database. *Nucleic Acids Research* (2020), **48** D383–D388. doi:10.1093/nar/gkz827.
